# Non-invasive brain stimulation over the Frontopolar Cortex promotes willingness to exert cognitive effort in a foraging-like sequential choice task

**DOI:** 10.1101/2024.09.27.615545

**Authors:** Mario Bogdanov, Laura A. Bustamante, Sean Devine, Signy Sheldon, A. Ross Otto

**Author notes:** Corresponding author: Mario Bogdanov, PhD McLean Hospital Center for Depression, Anxiety, and Stress Research 115 Mill Street Belmont MA 02478.

## Abstract

Individuals avoid spending cognitive effort unless expected rewards offset the perceived costs. Recent work employing tasks that provide explicit information about demands and incentives, suggests causal involvement of the Frontopolar Cortex (FPC) in effort-based decision-making. Using transcranial direct current stimulation (tDCS), we examined whether the FPC’s role in motivating effort generalizes to sequential choice problems in which task demand and reward rates vary indirectly and as a function of experience. In a double-blind, within-subject design, 46 participants received anodal (i.e., excitatory) or sham stimulation over the right FPC during an Effort Foraging Task, which required choosing between harvesting patches for successively decreasing resources or traveling to replenished patches by performing a cognitive task with environment-specific difficulty. As expected, participants exited patches later (i.e., displayed lower exit thresholds) when travelling required greater (versus less) effort, indicating increased travel costs in high-effort environments. Under anodal tDCS, the difference in exit thresholds between environments was significantly smaller relative to sham. Finally, individual differences analyses hint that participants with lower self-reported motivation to exert effort exhibited greater travel cost reductions following tDCS. Together, these findings support the theorized causal role of the FPC in motivating cognitively effortful behavior, expand its role to more ecologically valid serial decisions and highlight the potential for tDCS as a tool to increase motivation with potential clinical applications.

**Significance statement:** Uncovering the neural mechanisms regulating engagement in effortful behavior is crucial, as it will improve our understanding and treatment of conditions characterized by reduced motivation, e.g., apathy and anhedonia. The Frontopolar Cortex (FPC) has been implicated in increasing effort exertion in settings that provide explicit information about effort demand and reward. Using transcranial direct current stimulation (tDCS), we investigated whether the FPC retains its motivating capacity in sequential choice problems that vary effort and reward indirectly. We demonstrate that FPC stimulation decreases cognitive effort-based travel costs in an Effort Foraging Task, indicating a causal and general involvement of the FPC in motivating effortful behavior, highlighting the potential of tDCS as a new avenue for increasing motivation with potential clinical applications.

## Introduction

A growing body of theoretical and experimental work suggests that individuals’ decisions to engage in cognitively demanding behavior are (partly) determined by a cost-benefit trade-off that gauges the subjective value of expected outcomes based on the effort required to obtain them (Kool & Botvinick, 2018; Shenhav et al., 2017; Westbrook & Braver, 2015). This neuroeconomic perspective on effort-based decision-making has gained substantial empirical support: behavioral experiments have reliably demonstrated participants’ tendency to avoid spending effort and to discount prospective rewards with increasing task demands (Bogdanov et al., 2021; Devine & Otto, 2022; Kool et al., 2010; Westbrook et al., 2013), while neuroimaging studies linked these processes to brain structures implicated in reward processing and cognitive control allocation, including the ventral striatum (VS), ventromedial prefrontal cortex (vmPFC), dorsal anterior cingulate cortex (dACC), and frontopolar cortex (FPC) (Botvinick et al., 2009; Chong et al., 2017; Lopez-Gamundi et al., 2021; Vassena et al., 2014; Westbrook et al., 2019).

Although these findings provided valuable insights into the neural systems associated with effort-based decision-making, they do not allow to infer causality due to the correlational nature of common neuroimaging approaches (Krakauer et al., 2017). However, establishing causal brain-behavior relationships – and developing interventions to increase effort exertion – is critical, given growing evidence suggesting that reduced willingness to exert cognitive effort may contribute to transdiagnostic symptoms of apathy and anhedonia (Husain & Roiser, 2018; Jurgelis et al., 2021; Le Bouc et al., 2023).

To more directly investigate how modulations of cortical activity affect behavior, researchers have started to employ non-invasive brain stimulation techniques (Bogdanov et al., 2017, 2018; Bogdanov & Schwabe, 2016; Fregni et al., 2005; Polanía et al., 2018). Notably, a recent study demonstrated the efficacy of anodal (excitatory) compared to cathodal (inhibitory) or sham transcranial direct current stimulation (tDCS) over the right FPC for boosting willingness to expend physical and cognitive effort, suggesting domain-general involvement of this structure in effort valuation and decision-making (Soutschek et al., 2018). So far, this has only been tested using discounting paradigms that are frequently employed to study neural mechanisms of effort-based decision-making and typically involve choices between two explicitly described options—exerting more (versus less) effort for larger (versus smaller) rewards—in discrete trials where effort exertion is often hypothetical or occurs substantially delayed from the time of choice. Yet, recent evidence suggests that people may also gradually and momentarily adjust application of (effortful) cognitive control according to experienced fluctuations in the average reward rate of the current environment (Lin et al., 2022; Otto & Daw, 2019). Thus, if the FPC is truly instrumental in motivating effort across domains and choice context, experimentally modulating its activity should also affect effort exertion in tasks involving sequential decisions and variations in environmental richness.

To investigate this hypothesis, we conducted a double-blind, within-subject experiment in which participants underwent anodal and sham tDCS over the right FPC while completing the recently developed cognitive Effort Foraging Task (EFT; Bustamante et al., 2023). The study of foraging serial decision problems originated in ethology, valued for its strong theoretical foundations and ecological validity (Stephens & Krebs, 1986). Increasingly popular in neuroscience, foraging paradigms permit examination of how organisms estimate and balance time-varying reward rates and costs (Mobbs et al., 2018). These tasks have been instrumental in understanding decision-making and its neural mechanisms across species, including rodents (Carter & Redish, 2016; Kane et al., 2022), non-human primates (Hayden et al., 2011), and humans (Constantino & Daw, 2015; Hills et al., 2008, 2012; Kolling et al., 2012; Le Heron et al., 2020). The EFT is a patch foraging paradigm designed to quantify costs associated with cognitive effort, in which participants choose between repeatedly harvesting virtual patches for diminishing rewards versus performing a cognitively demanding task in order to travel to a new, replenished patch, adjusting effort exertion in accordance with experienced demand and reward levels. Consistent with the assumption that effort is treated as costly, participants, on average, exhibit significantly lower exit thresholds—that is, greater willingness to accept diminishing rewards to avoid effortful travel between patches—in high- versus low-effort environments (Bustamante et al., 2023, 2024). Further, these effort costs can be quantified by a foraging theory derived model, allowing to quantify that cost in terms of rewards (i.e., money) and to distinguish it from other important decision costs related to task performance, such as error aversion (Fleming et al., 2023).

Based on previous work linking FPC stimulation to increased motivation (Soutschek et al., 2018), we expected that anodal—relative to sham—tDCS would decrease effort costs associated with travelling, diminishing the difference in exit thresholds between effort environments and thus participants’ effort avoidance. Given the importance of dissociating error avoidance from effort costs (Fleming et al. 2023), we also controlled for participants’ travel task performance in our statistical approach. Further, we expected the stimulation to specifically affect exit thresholds, not participants’ cognitive task performance. Finally, in an exploratory analysis, we examined whether individual differences in working memory or self-reported tendency to exert cognitive effort (i.e., need for cognition) modulate to which degree tDCS affects foraging choice behavior.

## Materials and Methods

### Participants and experimental design

We recruited a total of 50 volunteers from the McGill community (40 female, 9 male, 1 ‘neither/other’; age range: 18 – 32 years; mean ± SD: 21.16 ± 2.41 years). Our original planned sample size was 40 participants, determined jointly based on earlier tDCS work and the consideration that the within-subjects manipulation employed here would yield greater statistical power than between-subject designs previously used in comparable tDCS studies (Bogdanov et al., 2017; Bogdanov & Schwabe, 2016; Ohmann et al., 2018; Soutschek et al., 2018, 2022). On the basis of pilot testing, we opted to recruit 10 additional participants to allow for potential issues related to the stimulation (e.g., technical errors, participants’ tolerance of the stimulation etc.). Five participants had to be excluded due to issues with the tDCS equipment for one (3 participants) or both (1 participant) of their testing sessions, leaving a final sample of 46 participants (36 female, 8 male, 1 ‘neither/other’; age range: 18 – 32 years; mean ± *SD*: 21.17 ± 2.50 years; see supplementary table S1). During recruitment, volunteers were excluded from participation if they met one or more of the following criteria (based on self-report): acute illnesses, lifetime history of any psychiatric or neurological disorder, or any contraindications specific to tDCS (i.e., family history of epilepsy, present or past head injuries, metal implants in the head, pacemakers etc.). Eligible participants provided informed consent and received either course credit or $45.00 CAD for completing both visits to the lab, with an additional (fixed) $5 bonus from the task. The study protocol followed the guidelines of the Declaration of Helsinki and was approved by the McGill Research Ethics Board.

We used a double-blind, sham-controlled within-subject design, in which participants completed a cognitive Effort Foraging Task (EFT, see below) under both anodal (active) and sham stimulation of the FPC. We opted against including a cathodal stimulation condition as previous work suggests it had no effect on participants’ willingness to exert effort (Soutschek et al., 2018). The order in which tDCS conditions were applied was counterbalanced between participants.

### Transcranial direct current stimulation

tDCS was administered using a Neuroconn Stimulator (Neuroconn, Germany). Electrodes were placed using rubber straps and an EEG cap to specify locations according to the standard 10-20 system. Following Soutschek and colleagues (2018), a smaller electrode (5 × 5cm) was attached over the right FPC with its center over channel position Fp2. The second, larger electrode (5 × 7cm), which served as a reference, was placed over the vertex, with its center at channel position Cz (see Figure 1A). In line with our past work (Bogdanov et al., 2017; Bogdanov & Schwabe, 2016), we applied a current of 1.075mA during active tDCS, which has been shown to be very effective in modulating cognitive processes (Hoy et al., 2013). Given the different electrode sizes, this led to current densities of 0.043mA/cm^2^ and 0.031mA/cm^2^ over the FPC and vertex, respectively (see Figure 1B). The lower current density of the reference electrode was intended to reduce the likelihood that observed effects are due to a (cathodal) stimulation of the vertex instead of the intended enhanced activity in the FPC by anodal tDCS.

**Figure 1.**
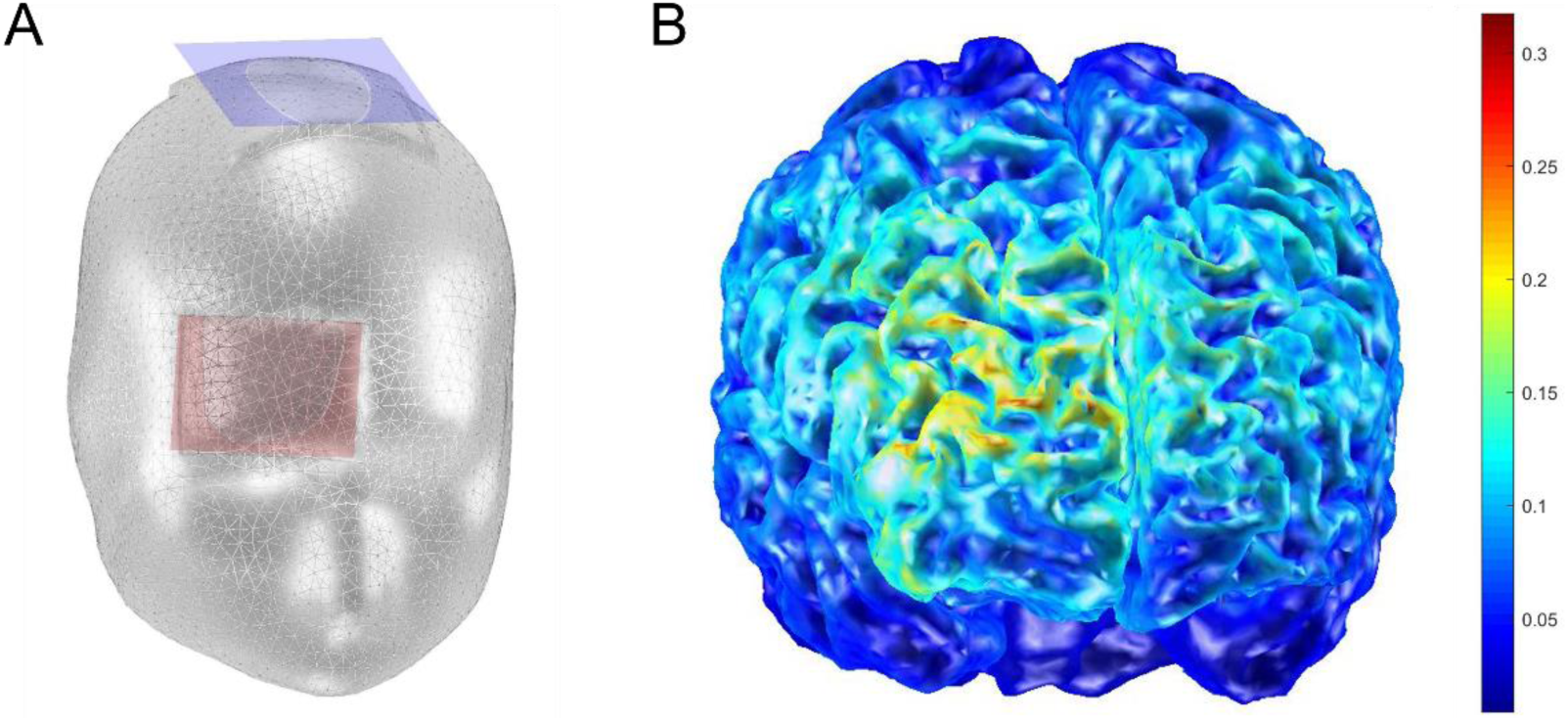
Transcranial direct current stimulation. (A) The active electrode (5x5cm, in red) was placed over the right FPC whereas the reference electrode (7x5cm, in blue) was placed over the vertex. (B) Simulation of the current density using the Comets2 toolbox for Matlab (Jung et al., 2013).

The electrode setup was identical in anodal and sham sessions. In both conditions, we applied current using an eight-second fade-in and a five-second fade-out window during which the current gradually ramped up and down, respectively (Bogdanov et al., 2017; Bogdanov & Schwabe, 2016). In the sham condition, fade-in was immediately followed by fade-out and no further stimulation took place after that. This was done to elicit the same initial sensation of tDCS in both conditions, making it difficult for participants to decode which condition they were in. Double blinding was achieved using preprogrammed codes provided by the stimulation system. For safety purposes, active tDCS was delivered for a maximum of 30 minutes after which the stimulator automatically initiated the fade-out and terminated the stimulation.

However, in the anodal tDCS condition, all participants received stimulation for the full duration of the EFT.

### Effort Foraging Task

We examined effort-based decision-making using a recently developed cognitive Effort Foraging Task (EFT; complete methods in Bustamante et al., 2023, Experiment 1), which was adapted from patch-foraging paradigms that have been shown to capture cost-benefit choice behavior aimed at optimizing reward rates in sequential decision tasks in an ecologically valid manner across species (Constantino & Daw, 2015; Kane et al., 2022; Mobbs et al., 2018) and that have been linked to FPC activity (Boorman et al., 2009; Daw et al., 2006).

In the EFT (see Figure 2), participants are placed in an environment representing a virtual ‘orchard’ where they could exploit patches (i.e., trees) for rewards (i.e., apples) by pressing a button on the keyboard, which in turn would earn them a monetary bonus at the end of the task. Trees could be harvested multiple times, but the number of received apples would decrease at a (mean) depletion rate of 0.88 with each consecutive harvest decision. Participants were required to harvest a tree at least once but were then free to decide to leave (i.e., ‘exit’) the current tree and travel to a new, replenished tree at any given time.

**Figure 2.**
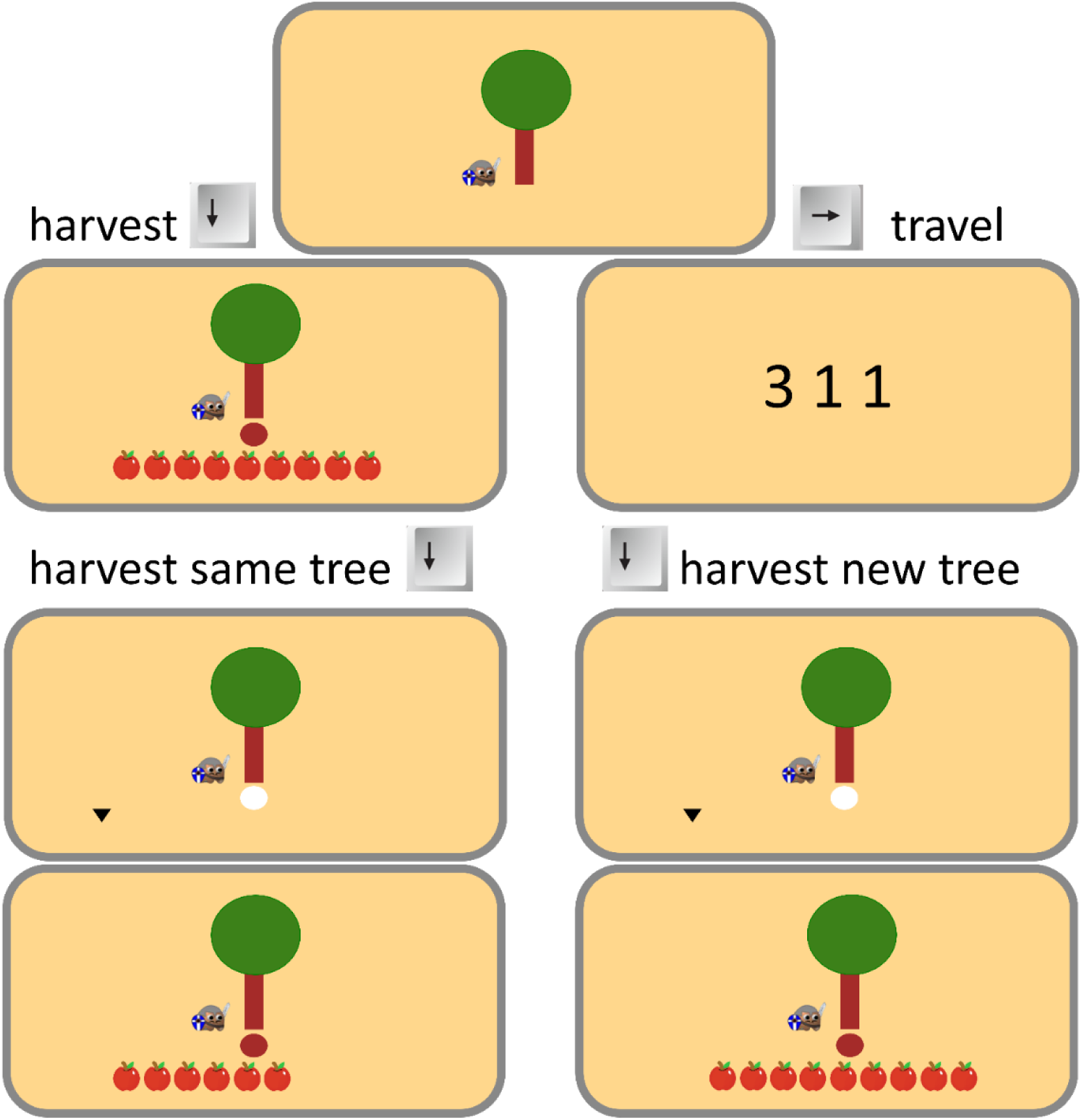
The cognitive Effort Foraging Task (EFT). Participants are placed in an orchard in which they can harvest trees for apples, i.e., rewards. In each trial, participants choose between harvesting the current tree or leaving for a different tree. Continuous harvesting decreases the number of apples received in each trial. Traveling leads to a replenished tree but requires completion of congruent (low-effort environments) or interference (high-effort environments) MSIT trials. The Effort level of the current environment (‘orchard’) was signaled by the background color: violet (low effort) or orange (high effort, depicted here).

To manipulate degrees of cognitive effort demand, participants were required to perform 6 trials of the Multi-Source Interference Task (MSIT; Bush & Shin, 2006) every time they chose to travel to a new tree, which effectively imposed a travel cost for exiting the current tree. In the MSIT, participants are presented with a display of three digits (range 0 - 3), with one digit being different from the other two (i.e., the ‘oddball’), and are required to press the key (‘1’, ‘2’, or ‘3’ on the keyboard) corresponding to the identity of the ‘oddball’ digit. In the low-effort condition, participants performed only congruent trials, in which the position of the oddball digit matched its identity, and the other digits were zeroes (e.g., participants had to press ‘1’ when the digits on display were 1-0-0, or ‘2’ when the digits were 0-2-0). In the high-effort condition, participants performed exclusively interference trials, in which location and identity of the oddball digit did not match, and the oddball was accompanied by digits other than zero (e.g., they had to press ‘3’ when the digits were 3-1-1).

We manipulated the environment (or orchard) effort level in a block-wise fashion—that is, every exit decision in a single block would lead to the same MSIT trial type. Participants completed two low-effort blocks and two high-effort blocks in counterbalanced order, each lasting four minutes, for a total of 16 minutes. The effort level of the current orchard was signaled by the background color of the screen (purple = low effort, orange = high effort). The foraging environment was held constant across blocks. Further, travel time between patches was fixed (8.33 seconds) across all environments, independent of participants’ MSIT performance. This was done in order to avoid the concern of unequal opportunity costs of time, which would be introduced by variable travel time durations and could have obfuscated the effects of effort on travel choices.

Following the original formulation of the EFT (Bustamante et al., 2023), participants were not required to perform the MSIT trials correctly in order to arrive at the next tree, nor were participants’ bonus earnings influenced by their MSIT task performance. However, in the practice we used a procedure to encourage participants to try their best at the travel task. In this procedure, participants had to complete 5 travel-length mini-blocks of MSIT trials successfully for both congruent and incongruent trial types. Successful completion was determined by not seeing the error feedback across the 6 trials, the error feedback was shown when participants made two errors in a row (including omission errors). This same feedback was implemented in the travel task used in the main phase of the EFT.

Earlier work has shown that choice behavior in patch foraging tasks can be formally characterized using the Marginal Value Theorem (MVT; Charnov, 1976), which prescribes that in order to optimize reward, an individual should estimate the average reward rate of the current environment and exit a tree as soon as the instantaneous reward provided by continued harvesting of a given patch falls below this average (Constantino & Daw, 2015; Kane et al., 2022). Critically, the elevated effort costs in the high-effort orchards are thought to decrease participants’ estimate of the average reward rate, leading to over-harvesting of trees and a lower exit threshold (i.e., the number of expected apples received for the next harvest at the time participants choose to exit the current tree) (Bustamante et al., 2023, 2024). Accordingly, we interpret the difference in exit thresholds between low- and high-effort environments as the perceived (effort) cost associated with travel.

### Procedure

Testing took place across two separate sessions, exactly 7 days apart. On day one, participants provided informed consent and completed a battery of questionnaires measuring traits related to motivation and impulsivity as well as symptoms of apathy and anhedonia, including: the Behavioral Inhibition and Activation System scales (BIS/BAS; Carver & White, 1994), the short form of the UPPS-P Impulsive Behavior Scale (Cyders et al., 2014), the Need for Cognition Scale (NFC; Cacioppo et al., 1984), the Snaith-Hamilton Pleasure Scale (SHAPS; Snaith et al., 1995), and the Dimensional Apathy Scale (DAS; Radakovic & Abrahams, 2014). Apart from the stimulation condition, the remainder of the protocol was identical on both days of testing. Participants first completed an automated operation span (OSPAN) task (Unsworth et al., 2005) that is commonly used to measure working memory capacity and has previously been shown to explain interindividual variance in responsiveness to external manipulations of choice behavior in complex decision-making tasks (Otto et al., 2013). Scores for all questionnaires and the OSPAN are presented in Table S1 in the supplement. After receiving adequate practice with the EFT – including familiarization with harvesting and travelling in the virtual orchard, as well as interference and congruent MSIT trials – tDCS electrodes were attached to the participants’ heads. Shortly after the start of the stimulation, participants began completing the main phase of the EFT, which took them approximately 20 minutes. In addition, participants completed a second task that is outside the scope of this paper for roughly 10 minutes before the stimulation was turned off. At the end of their second visit, participants were fully debriefed and received their compensation.

### Stimulation tolerance and manipulation check

Overall, participants tolerated the tDCS procedure well. No one terminated their testing sessions and no adverse side effects were reported. All participants reported that they felt a tingling sensation at the location of the active electrode during the fade-in period that subsided over time. At the end of each testing session, we asked participants whether they believed they had received active or sham stimulation. Due to a technical issue, we only have valid responses for a subset of participants (day 1: n = 37, day 2: n = 36, both sessions: 34). To statistically test the effectiveness of our blinding procedure, we compared the proportions of participants’ correct guesses regarding the stimulation conditions, finding that participants were not able to distinguish between tDCS conditions on either testing session (session 1: χ^2^ = 1.25, p = .264; session 2: χ^2^ = 2.38, p = .123). Moreover, we observed no significant differences in accuracy for tDCS guesses between both sessions (paired *t*(33) = -0.24, *p* = .813).

### Data analysis

*Inferential statistics.* We estimated a series of mixed-effects regressions using the *lme4* package in *R* (Bates & Maechler, 2009; R Core Team, 2020). Specifically, we used a mixed-effects linear regression to predict participants’ exit thresholds (i.e., expected rewards in number of apples in trials in which participants chose to leave the current patch) as a function of effort level (i.e., high- versus low-conflict MSIT trials, coded as 0.5 and -0.5, respectively), tDCS condition (i.e., anodal versus sham, coded as 0.5 and -0.5, respectively), session number (i.e., first vs. second session, coded as -0.5 and 0.5, respectively) and all possible interaction terms (taken as fixed effects). Effort level, tDCS condition and their interaction were also taken as random effects over participants. To predict accuracy in the MSIT, we used a mixed-effect logistic regression with trial type (congruent versus interference, coded as 0.5 and -0.5, respectively), tDCS condition, and their interaction (taken as fixed effects) and tDCS condition as a participant-specific random effect. To analyze response times (RTs), we used mixed-effects linear regression to predict log-transformed RTs (on correct trials only) as a function of MSIT trial type, tDCS condition and their interaction as both fixed effects, as well as random effects taken over participants. All categorical variables were effect-coded (-0.5 for sham stimulation, low effort condition and interference MSIT trials, 0.5 for anodal stimulation, high effort condition and congruent MSIT trials) and all continuous predictor variables were centered within participants (z-scored). In all models, random intercepts were taken over participants. We sought to keep the random effect structures of our models maximal (Barr et al., 2013), but eliminated predictor variables from the random effects structure of models if their inclusion caused convergence issues. Regression specifications for all analyses are provided in the supplemental materials (see Table S2).

#### Model-based analysis

In addition to regression-based analyses, we estimated a hierarchical Bayesian logistic mixed-effects model based on the MVT to derive cognitive effort costs following Bustamante et al. (2023). The original model had 3 parameters relevant to the cognitive effort version of the task: an inverse temperature (β), a cost of travel in the low effort condition (*c*_*low*_), and a marginal change in cost of travel from the low to high effort condition (*c*_*high*_). To test the performance of our model, we compared it to two other variants. In the ‘stimulation effect’ model, our main model of interest for this study, we adapted the original model to include a stimulation session effect on all 3 model parameters (β_*stim*_, *c*_*low*_*stim*_, *c*_*high*_*stim*_), resulting in 6 parameters total. The stimulation effect was modeled as an offset from the mean parameter (added to the mean for anodal, subtracted for sham). The model also included a 6 x 6 covariance matrix. With this configuration, we predicted the stimulation effect parameter for cognitive effort cost (*c*_*high*_*stim*_) would be negative, indicating reduced cognitive effort costs (i.e., lower effort avoidance) under stimulation. We did not predict significant differences in the other parameters by stimulation. In a third model, we tested the effect of session number on the stimulation effect model by including a session effect on all three model parameters (β_*stim*_, *c*_*low*_*stim*_, *c*_*high*_*stim*_) to create a 9 parameter ‘stimulation and session effect’ model. We conducted model comparison using log posterior likelihoods for these 3 models using leave-one-out cross-validation and WAIC (Devine et al., 2023; Vehtari et al., 2020).

For all models, priors on mean group-level effects parameters were: β ∼ 𝒩(0,0.5), *c*_*low*_ ∼ 𝒩(0,10), and *c*_*high*_ ∼ 𝒩(0,10). Priors on the stimulation group-level effects parameters were: β_*stim*_ ∼ 𝒩(0,0.25), *c*_*low*_*stim*_ ∼ 𝒩(0,5), and *c*_*high*_*stim*_ ∼ 𝒩(0,5). Priors on random variance were assumed to be half-normal: β ∼ 𝒩(0,0.5), *c*_*low*_ ∼ 𝒩(0,10), and *c*_*high*_ ∼ 𝒩(0,10), β_*stim*_ ∼ 𝒩(0,0.25), *c*_*low*_*stim*_ ∼ 𝒩(0,5), and *c*_*high*_*stim*_ ∼ 𝒩(0,5). Priors used for the session group- and participant-level effects of session were identical to those used for the stimulation effects. The covariance matrix Σ was defined using a Lewandowski-Kurowicka-Joe distribution with shape 1 (Lewandowski et al., 2009). Participant-level (random effects) parameters and their group-level (fixed effects) distributions were estimated using Markov Chain Monte Carlo (MCMC) sampling implemented in Stan with the CmdStanR package (4,000 samples, 2,000 warm-up samples, 4 chains; Stan, 2021). To evaluate the posterior distributions of individual model parameters, we calculated the probability of direction (*pd*), which reflects the proportion of the distribution that falls above or below 0 and can be interpreted as a Bayesian analogue to the frequentist *p*-value (Makowski et al., 2019). Notably, however, a key advance of the Bayesian statistics framework is to move beyond binary cutoffs for statistical significance and instead focus on the likelihood of an effect falling within a certain range. Data and code for all analyses will be available at OSF at time of final publication.

## Results

### Anodal stimulation over the FPC reduces cognitive effort-related travel costs

We hypothesized that excitatory stimulation to the FPC would decrease participants’ cognitive effort cost (i.e., increase willingness to complete MSIT interference relative to congruent trials). We approached this in two ways (see Data Analysis section). Firstly, we ran a model-agnostic analysis of exit thresholds (i.e., the number of apples a participant expected to receive when they chose to exit a patch) using mixed-effects linear regression (coefficient estimates in Table 1). Secondly, we performed an MVT model-based analysis of changes in the cognitive effort cost parameter due to stimulation (model estimates in Table 2).

**Table 1.**
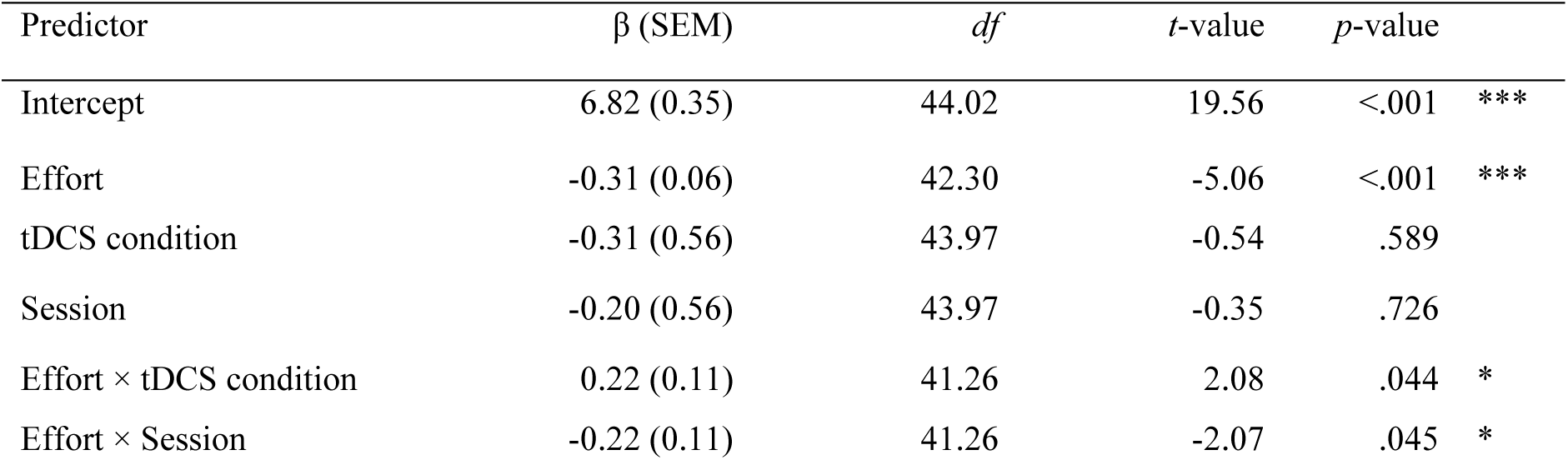
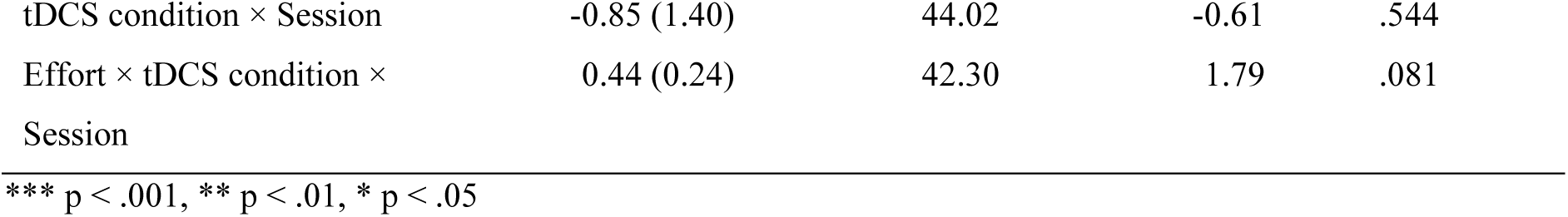
Mixed-effects regression model predicting exit thresholds by stimulation condition, effort level, and session number.

| Predictor | $\beta$ (SEM) | $df$ | $t$-value | $p$-value | |
| --- | --- | --- | --- | --- | --- |
| Intercept | 6.82 (0.35) | 44.02 | 19.56 | <.001 | *** |
| Effort | -0.31 (0.06) | 42.30 | -5.06 | <.001 | *** |
| tDCS condition | -0.31 (0.56) | 43.97 | -0.54 | .589 |  |
| Session | -0.20 (0.56) | 43.97 | -0.35 | .726 |  |
| Effort × tDCS condition | 0.22 (0.11) | 41.26 | 2.08 | .044 | * |
| Effort × Session | -0.22 (0.11) | 41.26 | -2.07 | .045 | * |

|  |  |  |  |  |
| --- | --- | --- | --- | --- |
| tDCS condition × Session | -0.85 (1.40) | 44.02 | -0.61 | .544 |
| Effort × tDCS condition ×<br>Session | 0.44 (0.24) | 42.30 | 1.79 | .081 |

\*\*\* $p < .001$ , \*\* $p < .01$ , \* $p < .05$

**Table 2.**
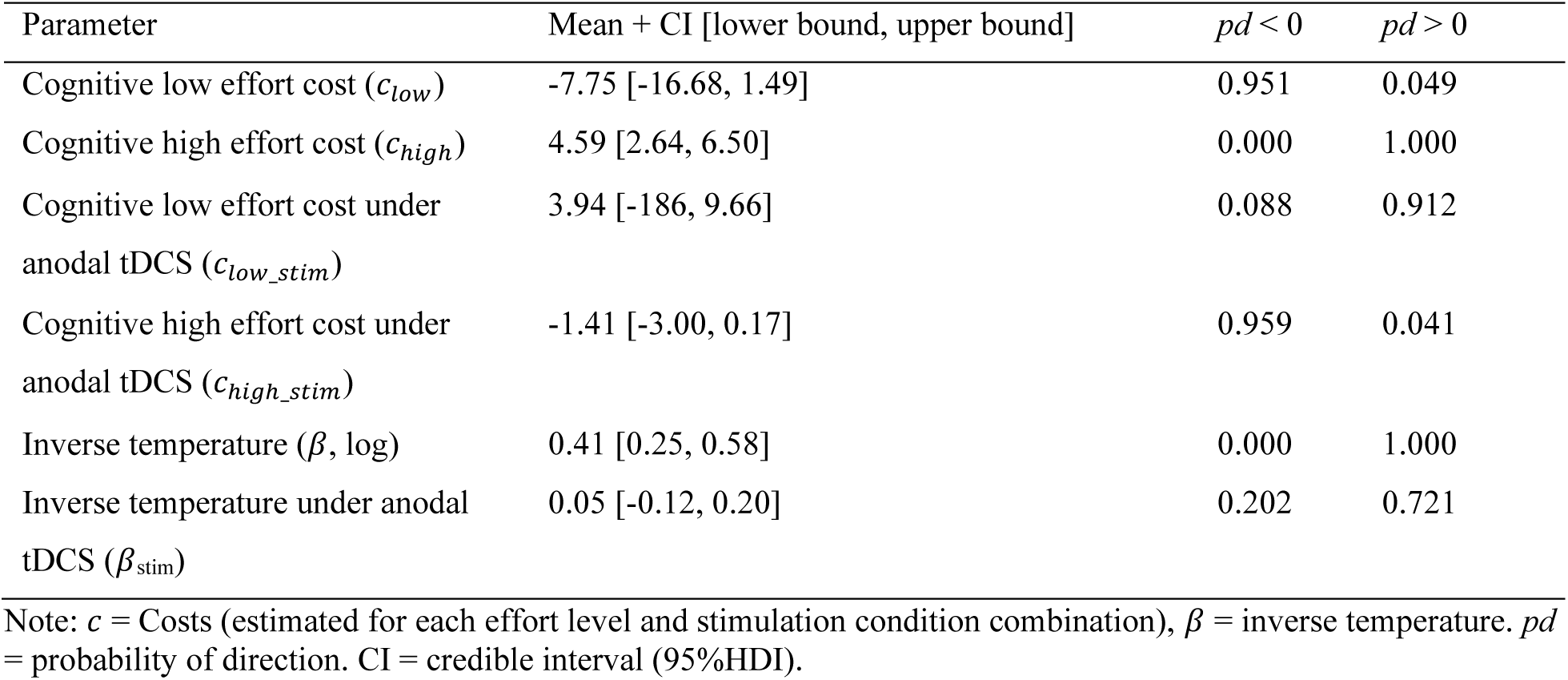
Group level posterior distributions of parameter estimates derived from the Hierarchical Bayesian Model based on the MVT.

| Parameter | Mean + CI [lower bound, upper bound] | $pd < 0$ | $pd > 0$ |
| --- | --- | --- | --- |
| Cognitive low effort cost ($c_{low}$) | -7.75 [-16.68, 1.49] | 0.951 | 0.049 |
| Cognitive high effort cost ($c_{high}$) | 4.59 [2.64, 6.50] | 0.000 | 1.000 |
| Cognitive low effort cost under anodal tDCS ($c_{low\_stim}$) | 3.94 [-186, 9.66] | 0.088 | 0.912 |
| Cognitive high effort cost under anodal tDCS ($c_{high\_stim}$) | -1.41 [-3.00, 0.17] | 0.959 | 0.041 |
| Inverse temperature ($\beta$, log) | 0.41 [0.25, 0.58] | 0.000 | 1.000 |
| Inverse temperature under anodal tDCS ($\beta_{stim}$) | 0.05 [-0.12, 0.20] | 0.202 | 0.721 |

Note: $c$ = Costs (estimated for each effort level and stimulation condition combination), $\beta$ = inverse temperature. $pd$ = probability of direction. CI = credible interval (95% HDI).

#### Model-agnostic analysis of exit thresholds

In the EFT, the longer a participant delays leaving the patch in the high- relative to low-effort environments, the larger their inferred effort cost. Delaying patch leaving indicates willingness to accept diminishing rewards in order to avoid the (effort) costs associated with traveling. Replicating previous work examining EFT behavior in a larger sample (Bustamante et al., 2023) we found that, overall, participants harvested trees longer before choosing to leave in high-effort, compared to the low-effort environments, as evidenced by a significantly lower exit threshold in high-effort environments (high effort: mean ± sem = 7.03 ± 0.37, low effort: mean ± sem = 7.31 ± 0.35, main effect of Effort: *p* <.001; Figure 3 and Table 1). We predicted that under anodal stimulation we would observe a smaller difference in exit threshold between effort levels, due to decreased perceived cost of traveling in the high- relative to the low-effort environment.

**Figure 3.**
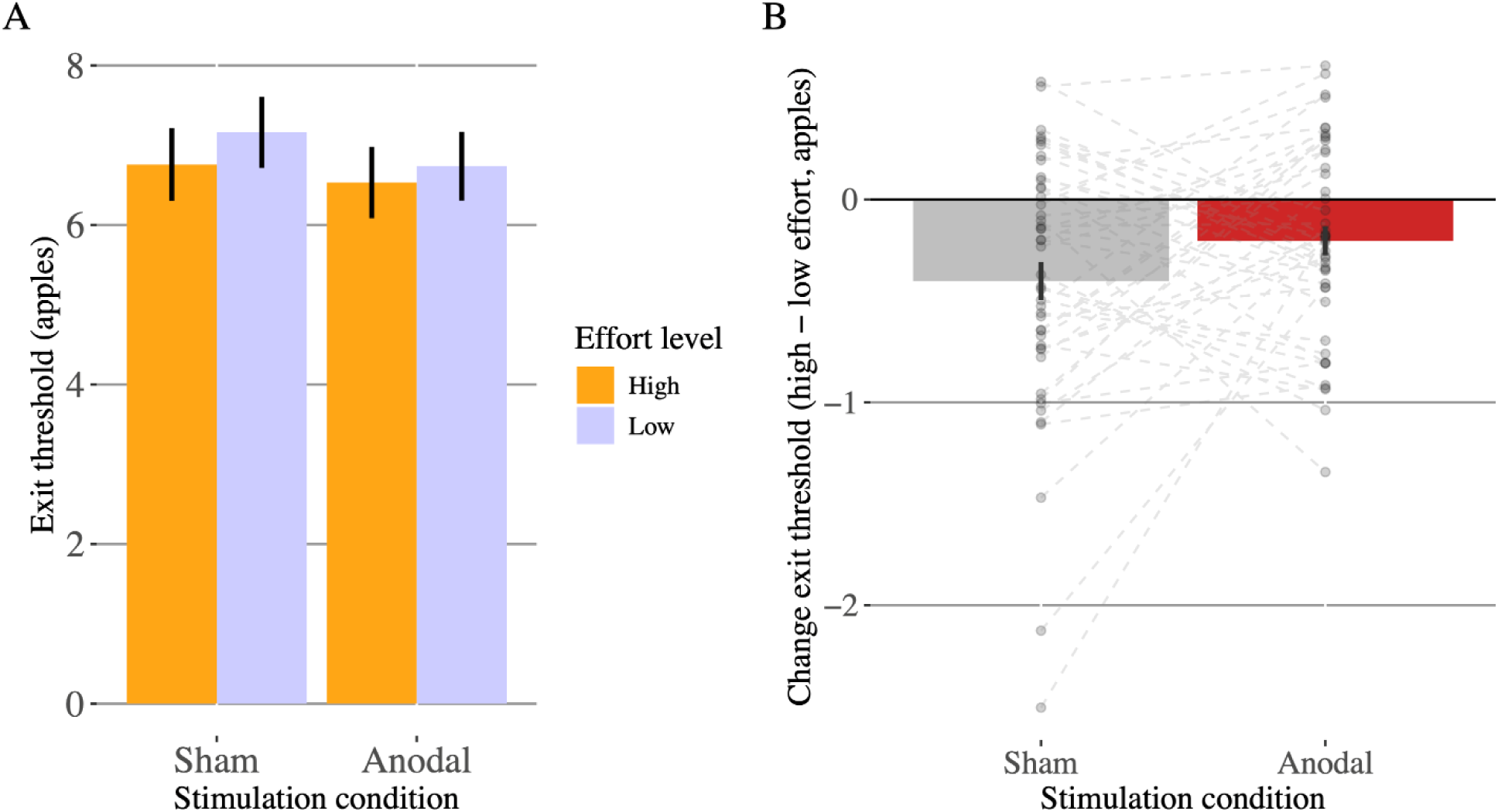
The effect of tDCS on participants’ exit thresholds. (A) Anodal stimulation over the FPC did not change the overall exit threshold at which participants chose to leave a given tree for either effort level in the MSIT. However, anodal stimulation significantly reduced the difference between high and low effort trials, suggesting a reduction in the relative effort-related travel costs in high-effort environments (B).

In line with our prediction, we found a significant interaction between environment effort level, and tDCS condition (*p* = .044), suggesting that anodal tDCS diminished the difference in exit thresholds between high and low-effort environments seen in the sham condition (see Figure 3). Importantly, this effect was specific to the change in exit threshold between conditions, as the overall exit thresholds in the task were not reliably affected by stimulation (main effect of tDCS condition: *p* = .589), indicating that tDCS did not change foraging behavior per se.

Given the within-subject, two-day design of our study, our regression model also included session number, which allowed us to control for practice effects. We observed a significant Effort × session interaction (*p* = .045), indicating the difference in exit thresholds between low- and high-effort environments was larger on the second day of testing. Importantly, however, we did not find any other significant effects of session (main effect of session: *p* = .726; tDCS × session: *p* = .544; effort × tDCS × session: *p* = .081), suggesting that the observed stimulation effects on foraging behavior did not depend on whether participants received anodal tDCS during their first or their second session.

#### MVT model-based analysis of exit thresholds

While the observed changes in average exit thresholds between low- and high-effort environments and tDCS conditions in the EFT can serve as proxy measures of how costly increased effort exertion is to participants, effort costs and stimulation effects can also be estimated more directly by fitting a computational model based on the MVT to the data, which has previously been shown to characterize behavior in patch-foraging tasks very well (Charnov, 1976; Constantino & Daw, 2015). We conducted model comparison using log posterior likelihoods for (1) a null model without specific stimulation effects, (2) a stimulation effect model, and (3) a model that accounts for both stimulation and session effects. All three models converged (R-hat < 1.01 for all parameters). The model comparison indicated that the stimulation effect model explained our data best (for more details, see supplement), therefore we will report model parameter estimates from this model only. Replicating previous findings by Bustamante et al. (2023), we found evidence for group-level cognitive effort avoidance expressed as a marginal increase in effort cost in units of approximately 4.6 apples in the high- compared to the low-effort environment (i.e, cognitive high effort cost (*chigh*) = 4.59 apples, *pd* > 0.001, see Table 2). Consistent with the model-agnostic results reported above, we found that high effort cost was reduced under anodal stimulation (group-level stimulation effect (*chigh_stim*) = -1.40 apples, *pd* < 0.041, see also Figure 4A), indicating increased willingness to exert cognitive effort under anodal tDCS. Stimulation did not affect cognitive low effort travel cost (*clow_stim* = 3.94, *pd* > 0.088) nor inverse temperature (*βstim* = 0.045, *pd* > 0.279), suggesting that tDCS increased motivation to engage in more effortful behavior specifically by altering how participants perceived the marginal increase in effort demands in high- versus low-effort environments. Examining individual participants (Figure 4B), we found that the mean posterior estimates for high-effort costs (c*high*) were mostly positive whereas those for the stimulation effect on high-effort costs (c*high_stim*) were mostly negative, indicating that, most individuals in our sample displayed effort avoidant behavior and that anodal tDCS mostly reduced high-effort costs across participants. However, it should be noted that the credible intervals around these participant-level estimates were (expectedly) large and partially crossed zero, constraining the strength of inferences that can be drawn from individual participant-level estimates.

**Figure 4.**
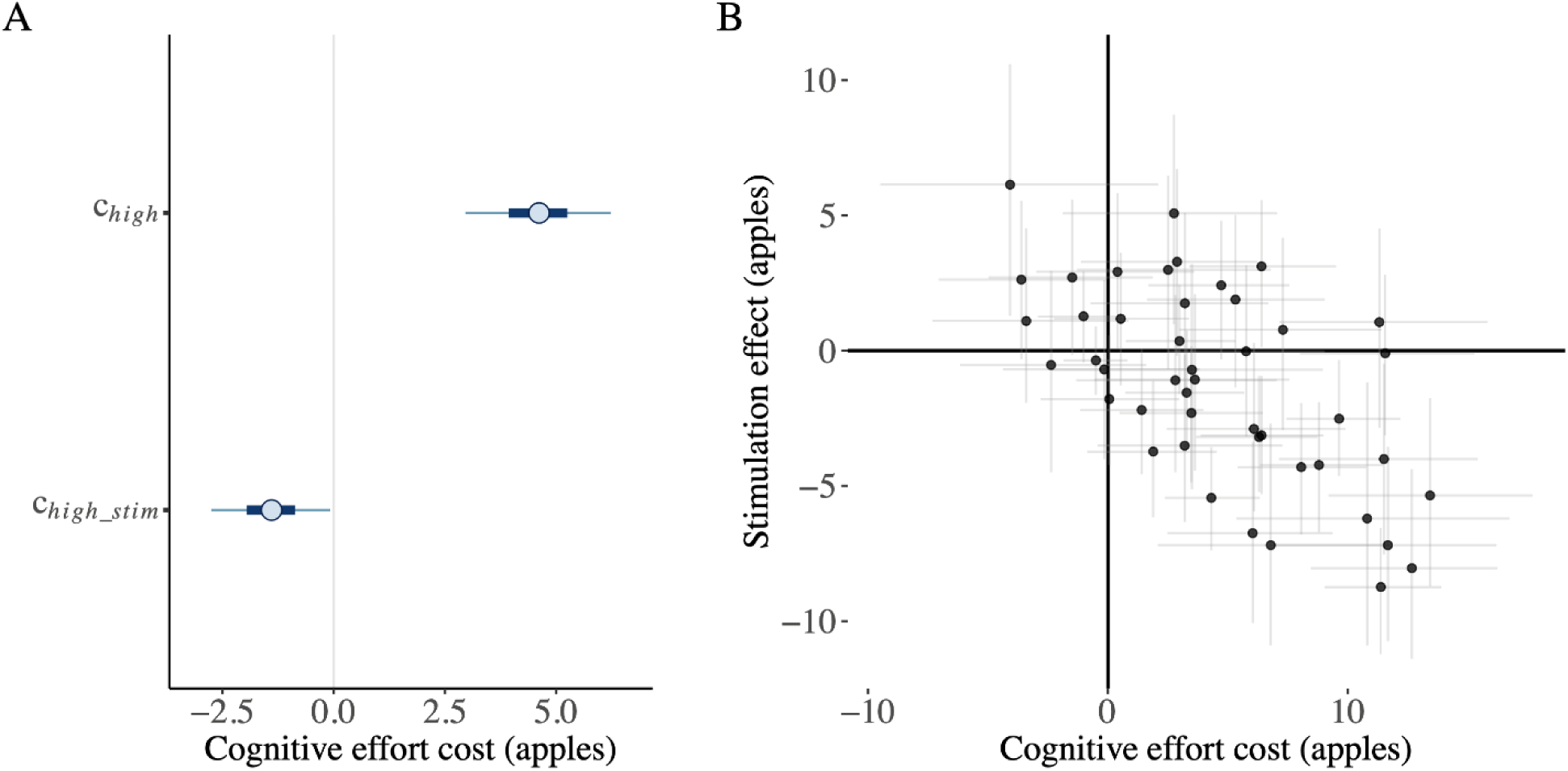
Posterior parameter estimates of high effort costs, and their modulation by anodal tDCS. (A) Group-level estimates from the MVT model suggest that high effort conditions are more costly than low effort conditions (c*high*), and a reduction of this effect under anodal tDCS (represented by negative value of c*high_stim*). (B) Individual estimates for each participant (i.e., the model’s random effects) indicate that all participants avoided effort and stimulation reduced effort avoidance, error bars indicate 80% highest density intervals.

### FPC stimulation did not affect performance in the MSIT

Examining performance in the MSIT using mixed-effects regressions (see Data Analysis section), participants responded faster and more accurately in congruent than interference trials, replicating previous work (Bush & Shin, 2006; Bustamante et al., 2023) and verifying successful manipulation of task demand (main effect of trial type for both measures: *p* < .001; see Table 3 and Figure 5). We also tested whether tDCS, beyond modulating effort-related travel cost sensitivity in the EFT, affected performance in the MSIT more directly, which could have possibly altered participants’ perception of travel costs (i.e. demands), in turn giving rise to the observed tDCS modulation of exit thresholds. Importantly, we did not find a main effect of tDCS condition on overall MSIT performance (accuracy: *p* = .302; log RTs: *p* = .372), nor an interaction between trial type and tDCS condition (accuracy: *p* = .466; log RTs: *p* = .127), indicating that performance did not differ meaningfully between brain stimulation conditions (for full coefficient estimates, see Table 3).

**Figure 5.**
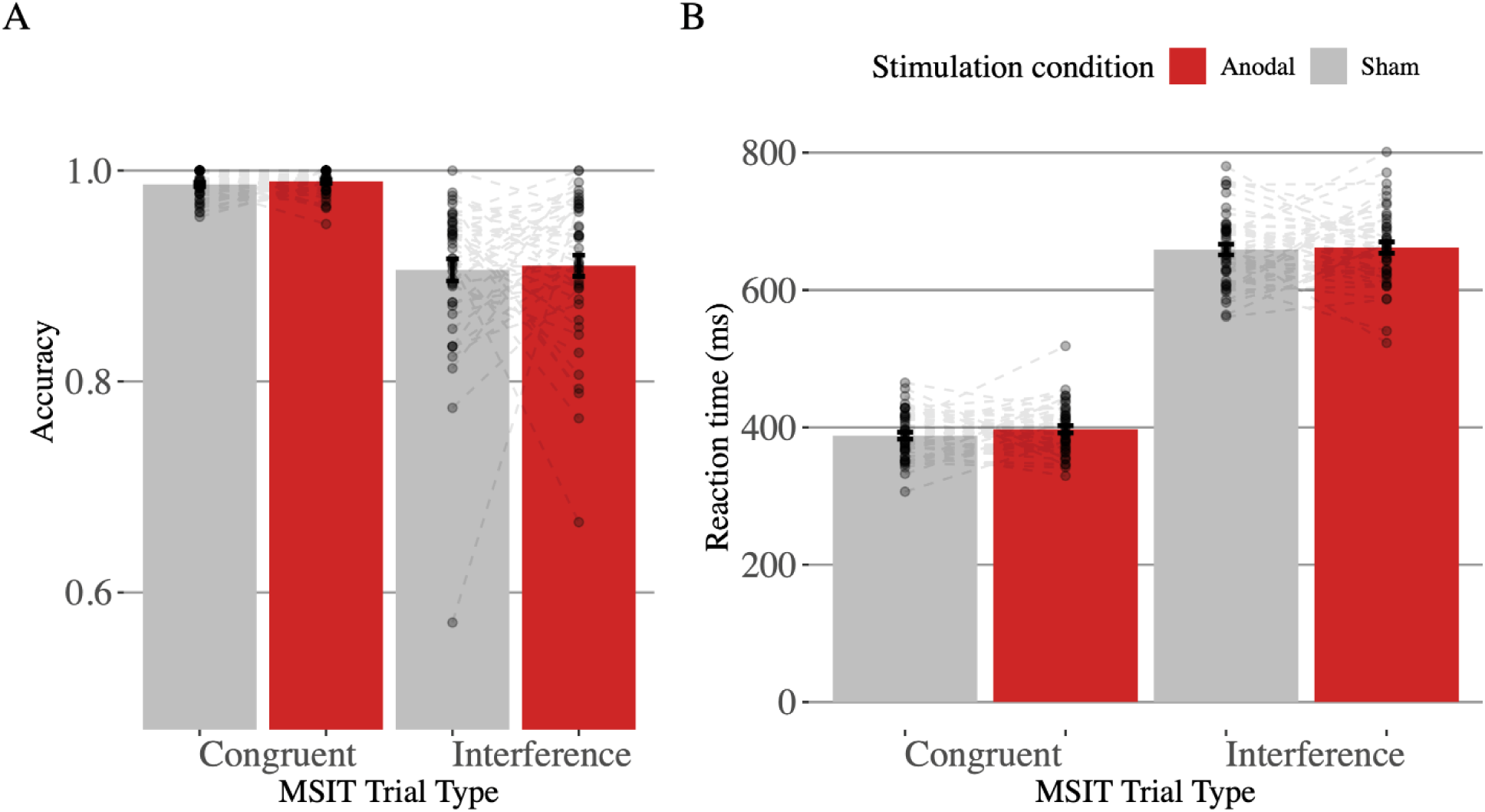
Performance in the MSIT. Participants were less accurate (A) and responded slower (B) in high effort compared to low effort MSIT trials. There was no effect of tDCS stimulation condition upon either performance measure.

**Table 3.** Results from a mixed-effects regression model predicting MSIT performance (accuracy and log- transformed correct RTs) by stimulation condition and effort level.

| Predictor | Accuracy |  |  |  | Response times |  |  |  |  |
| --- | --- | --- | --- | --- | --- | --- | --- | --- | --- |
| | $\beta$ (SEM) | $z$-value | $p$-value | | $\beta$ (SEM) | $df$ | $t$-value | $p$-value | |
| Intercept | 3.53<br>(0.09) | 41.01 | <.001 | *** | 6.22<br>(0.01) | 45.05 | 695.00 | <.001 | *** |
| Trial type<br>(congruent vs<br>interference) | 2.08<br>(0.09) | 22.39 | <.001 | *** | -0.52<br>(0.01) | 45.09 | -75.15 | <.001 | *** |
| tDCS condition | 0.18<br>(0.18) | 1.03 | .301 |  | 0.01<br>(0.02) | 44.91 | 0.90 | .372 |  |
| Trial type ×<br>tDCS condition | 0.14<br>(0.19) | 0.73 | .466 | | 0.02<br>(0.01) | 45.46 | 1.56 | .127 | |

\*\*\* $p < .001$ , \*\* $p < .01$ , \* $p < .05$

### Individual differences in MSIT performance did not affect exit thresholds

We also examined whether participants’ MSIT performance predicted (changes in) exit thresholds, and further, whether anodal tDCS would affect the relationship between MSIT performance and effort-related travel costs by estimating two additional regressions predicting exit thresholds which included MSIT accuracy and correct RTs (averaged separately for each combination of tDCS condition and effort level) as additional predictor variables (for full coefficient estimates, see Table 4). These analyses, in effect, address the possibility that participants’ subjective travel costs in the EFT might stem from MSIT performance rather than environment effort level—for example, participants who performed more accurately in the MSIT might have perceived travelling in the high-effort environment as less costly. By controlling for individual differences in task performance (i.e., MSIT accuracy and RTs in each experimental condition), we can directly examine the degree to which exit decisions were driven by participants’ effort aversion instead of—or in addition to—error aversion. Importantly, we found a significant main effect of MSIT accuracy upon exit thresholds (*p* < .001), indicating that participants who were more accurate (i.e., performed better) left trees earlier, and suggesting that they perceived travel to be smaller across both environment effort levels and both tDCS conditions. However, we did not observe significant interactions between MSIT accuracy and effort level (*p* = .368), nor between MSIT accuracy and tDCS condition (tDCS condition × accuracy: *p* = .455) suggesting that participants’ MSIT error rates did not affect participants’ subjective travel costs, and moreover, tDCS stimulation did not significantly modulate this relationship (Effort × tDCS condition × accuracy: *p* = .320). Critically, the main effect of environment effort level (*p* < .001) upon exit thresholds and the interaction of effort level and tDCS (accuracy: *p* = .024, RT: *p* = .043) remained significant, suggesting against the possibility that the reported stimulation effects on participants’ foraging behavior were caused by tDCS-induced changes in cognitive task performance. In other words, these control analyses indicate that tDCS modulated the impact of travel costs—instantiated here as environment effort level— upon exit thresholds over and above the potential contribution of MSIT error rates. To further support the notion that effort and error aversion are separable within the EFT, we conducted two additional analyses in which we correlated the model-derived high-effort costs (*chigh*; described above) with individuals’ accuracy and RTs in the MSIT (Bustamante et al., 2023, 2024). As expected, there was no significant association between effort costs and these performance metrics (accuracy: *r*(44) = 0.022, *p* = .886; RT: *r*(21) = 0.187, *p* = .394).

**Table 4.**
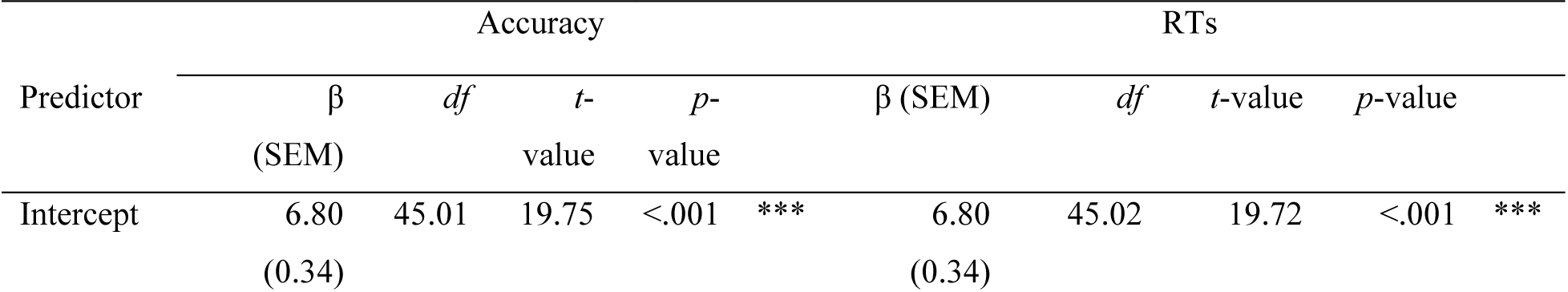
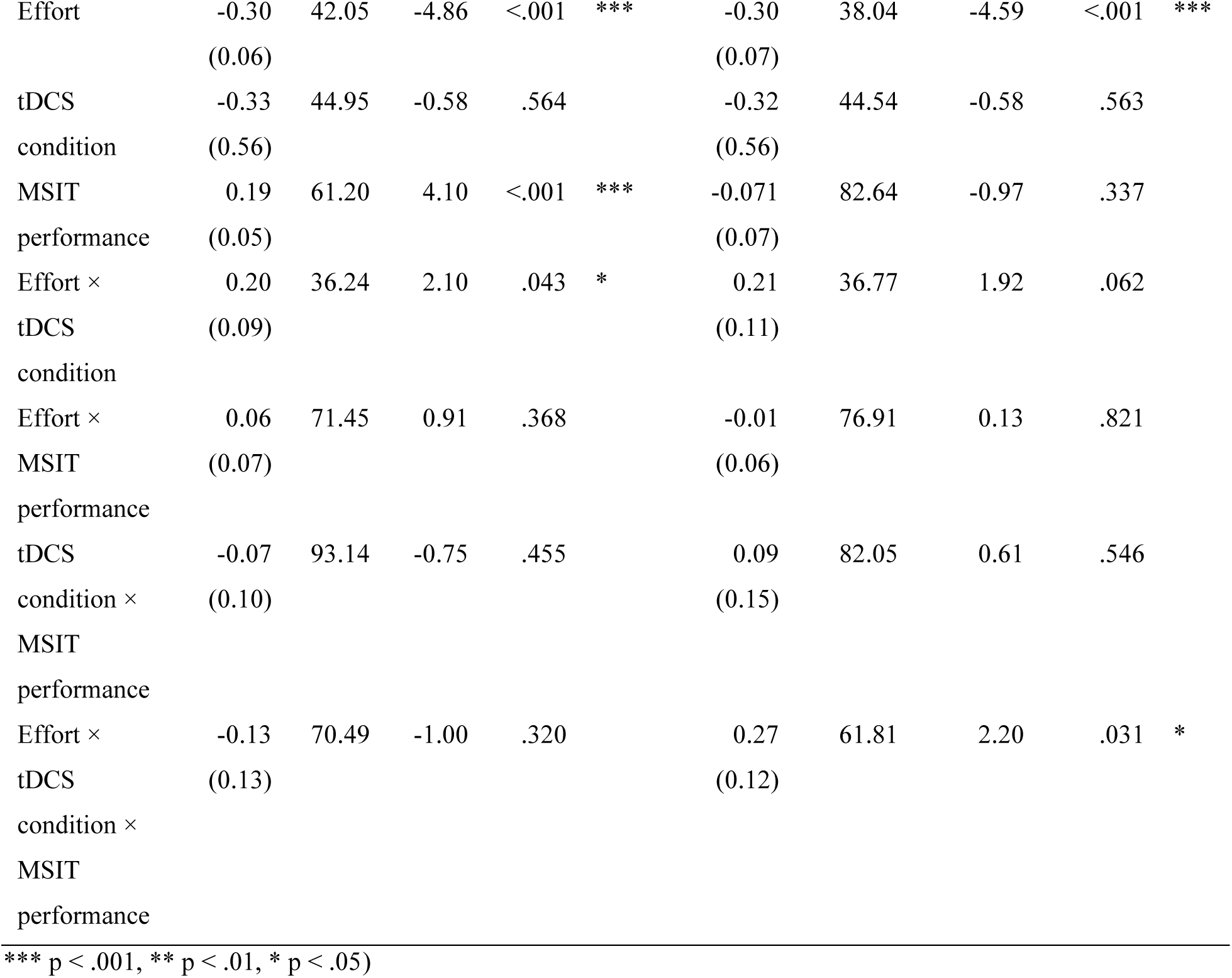
Results from a mixed-effects regression model predicting exit thresholds by stimulation condition, effort level, and task performance measure in the MSIT (i.e., accuracy or RTs, with separate averages for each combination of effort level and tDCS condition).

With respect to RTs, we did not observe a significant main effect of correct MSIT RTs on exit thresholds (*p* = .337) nor significant two-way interactions between RTs and effort level (*p* = .821) or tDCS condition (*p* = .546). However, we did observe a significant Effort × tDCS condition × RT interaction (*p* = .044; see Table 4), indicating that the effect of active stimulation on differences in exit thresholds between effort conditions was stronger for participants with slower correct MSIT RTs. While this effect was unexpected and should be interpreted with caution, this may suggest that anodal stimulation of the FPC may be particularly beneficial for individuals who perform slower in the MSIT.

<u>Individual differences in self-reported motivation to exert effort and Working Memory Capacity</u> Previous research has revealed a stronger preference to avoid cognitive effort in individuals with lower ‘Need for Cognition’ (NFC), a self-report trait thought to reflect an individual’s intrinsic motivation to engage in demanding mental activities (Cacioppo et al., 1984; Kührt et al., 2021; Sandra & Otto, 2018; Westbrook et al., 2013; Yan & Otto, 2020; Zerna et al., 2023; Zhang et al., 2022). Accordingly, in an exploratory analysis, in which we added participant-level NFC scores as a predictor to our original regression model, we examined the relationship between NFC scores and exit thresholds in the EFT as well as the extent to which low- versus high-NFC participants were differentially responsive to tDCS (see Figure 6). Neither the main effect of NFC (*p* = .304) nor the NFC × Effort level interaction (*p* = .395) were significant, indicating that participants’ NFC scores did not predict overall exit thresholds nor the difference in thresholds between high- and low-effort environments when collapsed across stimulation sessions. However, we found a significant three-way interaction between effort level, tDCS condition and NFC (*p* = .045; see Table 5), suggesting that the tDCS-induced decrease in exit threshold differences between effort levels was more prominent in lower-NFC participants. This may hint at the possibility that anodal tDCS over the FPC is more effective in individuals with lower intrinsic motivation to engage in cognitively effortful behavior.

**Figure 6.**
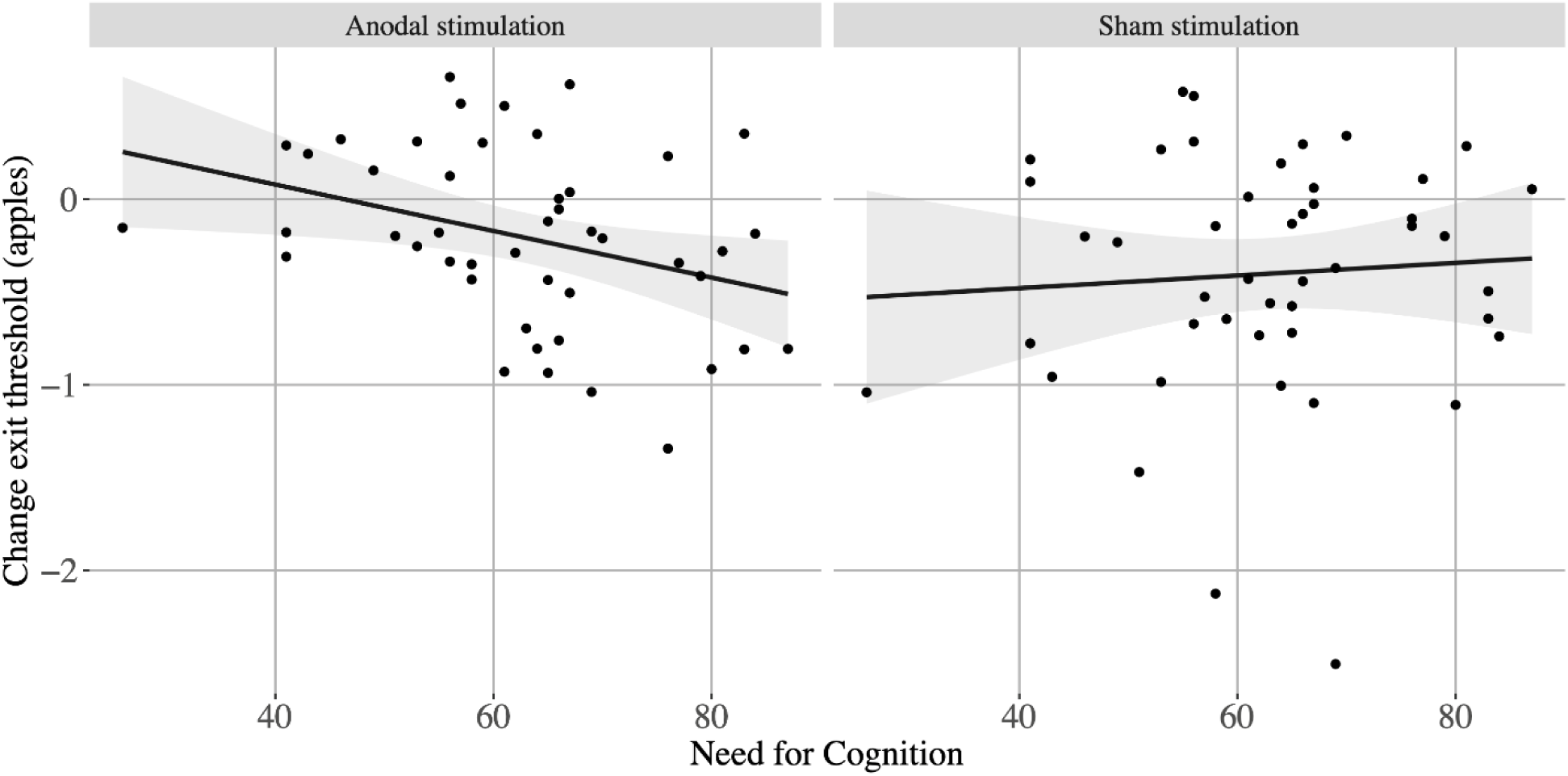
Predictive effects of Need for Cognition (NFC) scores on changes in exit threshold. More negative values in the change of exit thresholds indicate stronger effort avoidance, 0 indicates indifference between effort conditions. Under sham stimulation, there was no relationship between individuals’ NFC scores and the difference in exit thresholds between high- and low- effort conditions. In the anodal tDCS session, however, participants with lower NFC scores displayed smaller effort-related changes in exit thresholds, indicating a stronger reduction in effort-related travel costs during active stimulation of the FPC.

**Table 5.**
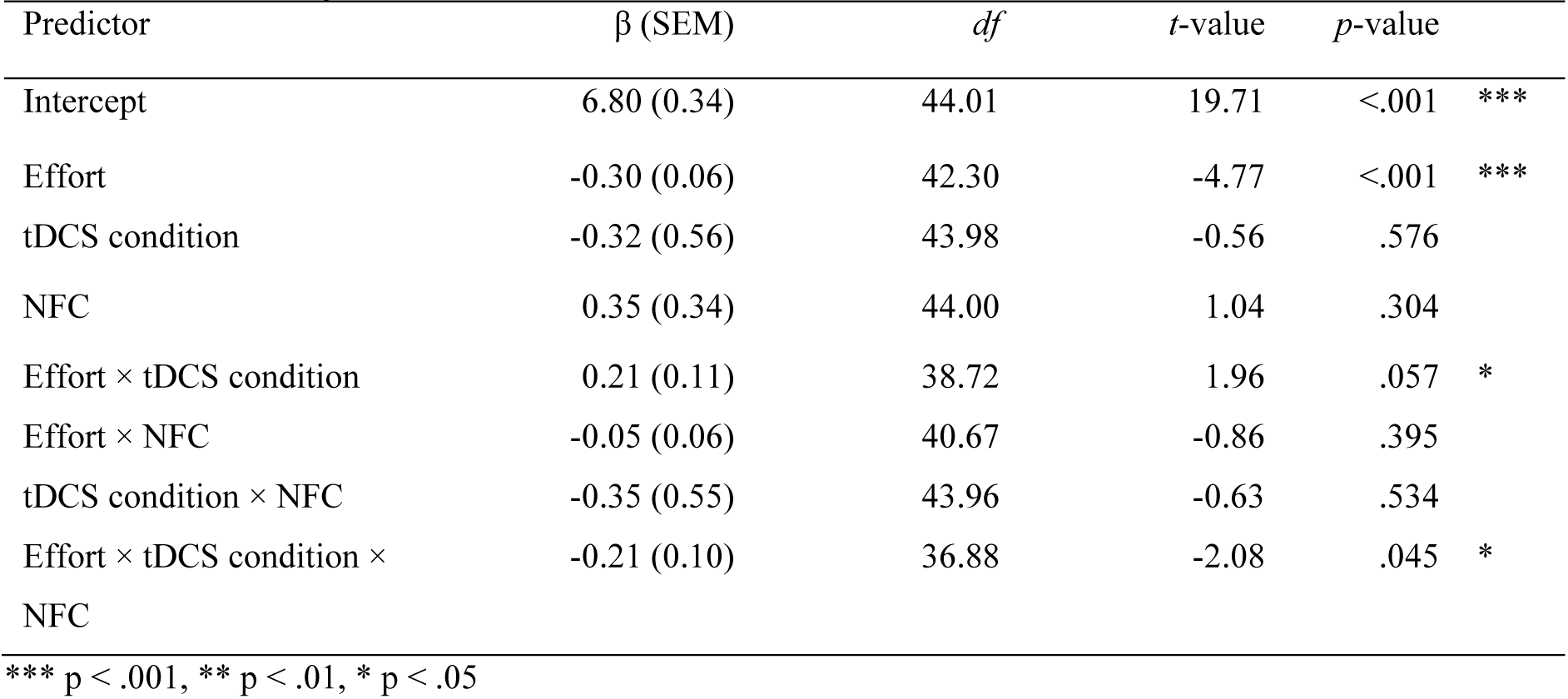
Results from a mixed-effects regression model predicting exit thresholds by stimulation condition, effort level, and Need for Cognition score.

| Predictor | $\beta$ (SEM) | $df$ | $t$-value | $p$-value | |
| --- | --- | --- | --- | --- | --- |
| Intercept | 6.80 (0.34) | 44.01 | 19.71 | <.001 | *** |
| Effort | -0.30 (0.06) | 42.30 | -4.77 | <.001 | *** |
| tDCS condition | -0.32 (0.56) | 43.98 | -0.56 | .576 |  |
| NFC | 0.35 (0.34) | 44.00 | 1.04 | .304 |  |
| Effort × tDCS condition | 0.21 (0.11) | 38.72 | 1.96 | .057 | * |
| Effort × NFC | -0.05 (0.06) | 40.67 | -0.86 | .395 | |
| tDCS condition × NFC | -0.35 (0.55) | 43.96 | -0.63 | .534 | |
| Effort × tDCS condition × NFC | -0.21 (0.10) | 36.88 | -2.08 | .045 | * |

\*\*\* $p < .001$ , \*\* $p < .01$ , \* $p < .05$

We were also interested in whether participants’ foraging behavior in the EFT might be related to individual differences in baseline working memory given the strong dependence of the MSIT upon executive functions (Bush & Shin, 2006; Fedorenko et al., 2011). To explore this possibility, we added participants’ session-specific OSPAN total scores—collected at baseline (i.e., before the tDCS administration) on each day of testing—to the mixed-linear regression predicting exit thresholds but found no significant main or interaction effects involving OSPAN scores upon exit thresholds (*p*s > .096; see Table S3 for full coefficient estimates).

Finally, we considered the possibility continued cognitive effort exertion in the EFT might have engendered fatigue, which could alter participants’ willingness to exert effort (Matthews et al., 2023; Müller et al., 2021), which could manifest as changes in foraging behavior over time—for example, increasing participants’ likelihood of making a harvest (versus travel) decision towards the end of a block. Accordingly, examined whether time-on-task may have affected exit thresholds and whether tDCS modulated these effects, by including (z-scored) exit decision number within each block as a proxy for time-on-task (and in turn, fatigue) as a predictor in our exit threshold-predicting regression. We observed no significant main effect of exit decision number within a block on exit thresholds (*p* = .318), nor any significant interaction effects involving within-block exit trials (all *p*s > .443), suggesting against an effect of fatigue on our findings (Table S4).

## Discussion

Identifying which brain regions are causally involved in guiding cognitive effort exertion is crucial to better understand—and ultimately treat—symptoms characterized by reduced motivation commonly observed across neurological and psychiatric conditions (Husain & Roiser, 2018). Recent tDCS work suggests a causal and potentially domain-general role for the FPC in increasing willingness to engage in cognitively demanding behavior, specifically in decision tasks that provide explicit information about effort costs and prospective rewards (Soutschek et al., 2018). However, if the FPC is indeed universally involved in effort-based choice behavior, this motivational effect should generalize to other decision contexts as well.

Thus, we here investigated how (excitatory) anodal—compared to sham—tDCS over the right FPC affects behavior in an Effort Foraging Task (Bustamante et al., 2023, 2024), which presents participants with sequential choice problems that rely on an indirect evaluation of effort costs based on the continuous experience of the average environmental reward rate. Replicating findings reported by Bustamante and colleagues (2023), under sham stimulation, participants chose to harvest trees for longer in high- compared to low-effort environments before leaving.

Importantly, however, anodal tDCS markedly reduced the difference in exit thresholds between environments, indicating increased willingness to exert cognitive effort. Presenting converging evidence from model-agnostic linear mixed-effects regression analyses and a hierarchical Bayesian model based on the MVT (Charnov, 1976), we demonstrate that excitatory stimulation of the FPC diminishes the subjective effort cost imposed by travelling in high-effort environments.

Our findings add to a growing body of work aimed at identifying the neural mechanisms that guide motivated behavior, corroborating compelling (but nevertheless correlational) neuroimaging evidence for the involvement of the FPC in motivating performance in incentivized, effortful tasks (Burke et al., 2013; Locke & Braver, 2008; Pochon et al., 2002), and demonstrating that the causal role of the FPC for effort-based decision-making is not limited to effort-discounting paradigms (Soutschek et al., 2018).

Effort-discounting tasks in particular are commonly used in the literature as they provide straightforward instantiations of effort-based choice, which assume that effort allocation decisions follow a cost-benefit analysis that balances expected effort costs and rewards (Shenhav et al., 2013, 2017; Westbrook & Braver, 2015). However, while they allow explicit, parametric manipulation of task demand and outcomes and have consistently produced convincing evidence that individuals generally tend to avoid effort exertion (Bogdanov, Renault, et al., 2022; Kool et al., 2010; Vogel et al., 2020; Westbrook et al., 2013), they only represent a subset of potential operationalizations of effort-based decision-making. To assess the generalizability of the FPC’s involvement in effort-based choice behavior, we employed a cognitive Effort Foraging Task (Bustamante et al., 2023). This task was built upon influential patch-foraging paradigms, but manipulates effort as the cost associated with travelling between patches in contrast to time, which is held constant (Constantino & Daw, 2015; Le Heron et al., 2020; Mobbs et al., 2018).

We chose the EFT as it possesses several desired properties for this experiment. First, foraging tasks capture sequential choice behavior in dynamic environments that may be more reflective of real-world decision-making and may not be well-characterized by effort-discounting paradigms (Carter et al., 2015; Carter & Redish, 2016; Gabay & Apps, 2021; Mobbs et al., 2018). This more indirect approach to assessing effort-based choices reduces the potential influence of demand characteristics on the experiment and could thus be more reflective of participants’ true preferences toward effort exertion (Bustamante et al., 2023). Second, whereas effort expenditure is often hypothetical or delayed in effort-discounting tasks, the EFT requires participants to exert cognitive effort immediately after travel choices, increasing its ecologically validity. Third, foraging behavior has been linked to FPC activity (Kolling et al., 2012; Le Heron et al., 2020; Mansouri et al., 2017), increasing the likelihood that behavior in the EFT would be sensitive to FPC stimulation. Fourth, choices in foraging tasks can be modeled according to the MVT (Charnov, 1976; Constantino & Daw, 2015), allowing us to formally quantify participant-specific effort costs and stimulation-induced changes in these costs. Fits of this computational model and formal comparisons against alternative models further demonstrate that participants indeed ascribed larger costs to travelling in the high- relative to low-effort environment, and that stimulation of the FPC decreased these cost estimates, supporting the conclusions drawn from the model-free analyses. Taken together, our findings thus not only conceptually replicate previous reports on the importance of the FPC in motivating effort exertion but demonstrate that this function expands beyond the specific context of effort-discounting paradigms. In addition, our study further highlights the suitability of the EFT as a measure of the malleability of (cognitive) motivation.

Statistically controlling for potential confounds, we found that stimulation effects on choice behavior were not modulated by the experimental session, which rules out influences of task familiarity on tDCS effects, nor by baseline working memory capacity. Notably, tDCS did not affect participants’ performance in the MSIT, which suggests against the possibility that stimulation-induced reductions in effort costs were brought about by changes in (subjective) task difficulty. Most importantly, we also aimed to examine whether individual differences in MSIT performance (i.e., accuracy and response times) affected foraging behavior. Given that interference (high-effort) MSIT trials, as intended, led to more erroneous responses, it is possible that our results could have reflected reduced error- instead of effort costs. Indeed, most paradigms used in effort-based decision-making conflate effort and difficulty, which renders drawing distinctions between these two constructs very difficult, as they are often strongly related (Dunn et al., 2019; Feghhi & Rosenbaum, 2021). Experimental approaches to address this concern, for example by using calibration procedures to balance error rates across demand conditions (Fleming et al., 2023), are nascent and the potential impact that these sort of individual calibration procedures have upon individuals’ subjective effort evaluation is not yet well understood. Thus, we instead opted to take a statistical approach to control for the contributions of error versus effort aversion to participants’ choices. Consistent with the idea that error costs might play an important part in effort avoidance, we found that accuracy predicted exit decisions in a general manner, but, importantly, this was not modulated by environmental demand level nor tDCS condition, suggesting that our findings cannot be fully explained by changes in error avoidance. This interpretation was further supported by the lack of correlation between incongruent trial accuracy and the effort costs parameter (*chigh*) from the computational model, suggesting again that these two phenomena are separable within the EFT. In sum, our findings indicate that the observed reduction in effort costs during anodal tDCS reflects a selective modulation of effort costs rather than cognitive performance or error avoidance.

We also investigated the possibility of that fatigue-like effects could be operating in the EFT which could have biased individuals to harvest trees for longer as a task block progresses, owing to a decreased willingness to exert cognitive effort (Jurgelis et al., 2021; Massar et al., 2018; Matthews et al., 2023) previously linked to the prefrontal cortex (Müller et al., 2021; Soutschek et al., 2022; Soutschek & Tobler, 2020). However, we found no evidence for a general effect of fatigue on foraging behavior nor fatigue effects specific to high-effort environments or tDCS condition. Finally, our exploratory analysis of participants’ NFC scores suggests that tDCS effects were stronger for individuals reporting lower Need for Cognition, a trait shown to be predictive of individuals’ effort evaluation (Otto et al., 2022; Sandra & Otto, 2018; Zerna et al., 2023; Zhang et al., 2022). This may suggest that stimulation might be most beneficial for people with comparatively low motivation and more pronounced effort aversion, consistent with similar baseline-dependent tDCS effects in other cognitive domains (Arciniega et al., 2018; Assecondi et al., 2021, 2022). However, the sample size in the current study, aimed to discover within-subject effects, does not permit conclusive judgments about these results. Thus, these exploratory findings should be interpreted with caution and warrant subsequent replication attempts in larger and more diverse samples, including older participants, as stimulation effects on the FPC may also differ by age (Soutschek et al., 2022).

While our findings support a causal role of the FPC in motivating cognitive effort exertion, we, importantly, do not propose that the observed effects solely depend on the FPC. Effort processing is thought to be served by a broad network of brain regions, including parts of the cognitive control network (e.g., lateral PFC and dACC) and the reward network (e.g., vmPFC or the VS), which are needed to perceive and estimate task demands and integrate them with reward information to form a choice signal indicating whether a potential course of action is worth the effort (Chong et al., 2017; Massar et al., 2015; Shenhav et al., 2013). Although tDCS should have primarily modulated excitability in the FPC, the stimulation possibly also affected distal brain areas and—in turn—choice behavior by downstream effects. For example, the FPC is reciprocally connected to the ACC, dlPFC, and the amygdala, it projects to the striatum (Chib et al., 2013; Hogeveen et al., 2022; Orr et al., 2015; Petrides & Pandya, 2007; Riedel et al., 2019), and is crucial for coordinating network activity across these structures (Ainsworth et al., 2022).

Additionally, prefrontal tDCS has shown to increase striatal dopamine levels (Bunai et al., 2021; Fonteneau et al., 2018; Fukai et al., 2019). Given the well-established importance of dopamine signaling for effort-based decision-making processes (Bogdanov, LoParco, et al., 2022; McGuigan et al., 2019; Salamone et al., 2016; Westbrook et al., 2020; Westbrook & Braver, 2016) and foraging behavior (Le Heron et al., 2020), tDCS possibly affected choice behavior by altering dopamine release. Taken together, our findings thus most likely result from a systemic rather than a focal modulation of neural activity.

While this study was conducted in young adults with no reported psychiatric disabilities, our findings may also have clinical implications. Compelling evidence suggests that altered effort processing may be a contributing factor of apathy and anhedonia, symptoms commonly observed across many neurological and psychiatric conditions, including major depression, schizophrenia, Parkinson’s disease and dementia (Culbreth et al., 2018; Husain & Roiser, 2018; Le Heron et al., 2018; McGuigan et al., 2019). First studies administering non-invasive brain stimulation to patients with treatment-resistant depression have demonstrated promising results in terms of reducing anhedonia (De Raedt et al., 2015; Fukuda et al., 2021; Rezaei et al., 2023; Sonmez et al., 2019), but most have targeted cortical regions outside the FPC. Our results suggest that anodal tDCS over the FPC, either on its own or complementary to psychotherapy (Nord et al., 2019), might be an effective, low-risk option to improve motivation in individuals experiencing apathy and anhedonia.

In conclusion, we provide first evidence for a causal role of the FPC in estimating the costs of effort exertion in sequential, foraging-like choice problems, which present a novel, ecologically valid alternative to the commonly employed explicit effort-discounting paradigms. We thereby corroborate and expand earlier findings on this area’s function in effort-based decision-making. These results enhance our understanding of the neural and computational mechanisms underlying motivation and cognition and may have useful implications in developing methods to increase engagement in cognitively demanding behavior in both clinical and non-clinical populations.

## Conflict of interest

The authors declare no competing financial interests.

### Acknowledgements

The research was supported by funding from the German Research Foundation (DFG, grant BO 5457/1-1), the Natural Sciences and Engineering Research Council of Canada (NSERC), the Canadian Foundation for Innovation (CFI), and the Centre for Research on Brain, Language and Music (CRBLM), the National Institute on Drug Abuse (NIDA, grant T32DA007261). We gratefully acknowledge the assistance of Marin Bergeron, Yiwei Cao, and Echo Wang in data collection and Nathaniel Daw for helpful discussions during the development of the computational model.

