## Supplemental material for "Non-invasive brain stimulation over the Frontopolar Cortex promotes willingness to exert cognitive effort in a foraging-like sequential choice task"

### Supplementary Methods and Materials

#### Demographic information

**Table S1**. Sample characteristics.

| | | Mean | Standard Deviation |
|---|---|---|---|
| **Age** | | 21.17 | 2.50 |
| **NFC** | | 62.65 | 13.16 |
| **BIS** | | 20.80 | 3.53 |
| **BAS** | | | |
| | **Drive** | 10.87 | 2.13 |
| | **Fun Seeking** | 12.13 | 1.97 |
| | **Reward Responsiveness** | 17.30 | 2.23 |
| **UPPS-P** | | | |
| | **Negative Urgency** | 11.37 | 2.06 |
| | **Positive Urgency** | 8.98 | 1.64 |
| | **Sensation Seeking** | 10.89 | 1.83 |
| | **Lack of Premeditation** | 9.70 | 1.52 |
| | **Lack of Perseverance** | 9.48 | 1.95 |
| **SHAPS** | | 1.37 | 2.87 |
| **DAS** | | | |
| | **Executive** | 10.37 | 4.54 |
| | **Emotional** | 7.70 | 3.69 |
| | **Initiation** | 8.35 | 3.77 |
| | **Overall** | 26.41 | 7.49 |
| **OSPAN** | | | |
| | **Total score on day 1** | 21.63 | 7.38 |
| | **Total score on day 2** | 25.78 | 4.60 |

NFC = Need for Cognition, BIS = Behavioral Inhibition System, BAS = Behavioral Activation System, UPPS-P = Impulsive Behavior Scale, SHAPS = Snaith-Hamilton Pleasure Scale, DAS = Dimensional Apathy Scale, OSPAN = Operation Span.

### Regression specifications for all analyses presented in the manuscript

**Table S2.** Regression model specifications.

| Analysis | Table in manuscript | Regression model |
|---|---|---|
| Exit thresholds predicted by stimulation condition,effort level, and session number | Table 1 | expected reward ~ 1 + effort level * tDCS condition * session + (1 + effort level * tDCS condition \| participant) |
| MSIT accuracy predicted by stimulation condition and effort level | Table 3 | correct response ~ trial type * tDCS condition + (1 + tDCS condition \| participant) |
| MSIT RTs predicted by stimulation condition and effort level | Table 3 | RTlog ~ trial type * tDCS condition + (1 + trial type * tDCS condition \| participant) |
| exit thresholds predicted by stimulation condition, effort level, and MSIT accuracy | Table 4 | expected reward ~ 1 + effort level * tDCS condition * MSIT accuracy + (1 + effort level * tDCS condition \| participant) |
| exit thresholds predicted by stimulation condition, effort level, and MSIT RTs | Table 4 | expected reward ~ 1 + effort level * tDCS condition * MSIT RTs + (1 + effort level * tDCS condition \| participant) |
| exit thresholds predicted by stimulation condition, effort level, and NFC score | Table 5 | expected reward ~ 1 + effort level * tDCS condition * NFC + (1 + effort level + tDCS condition \| participant) |
| exit thresholds by stimulation condition, effort level, and OSPAN performance | Table S3 | expected reward ~ 1 + effort level * tDCS condition * OSPAN total score + (1 + effort level * tDCS condition \| participant) |
| effort level, and within block exit trial number | Table S4 | expected reward ~ 1 + effort level * tDCS condition * exit trial number + (1 + effort level + tDCS condition + exit trial number + effort level:exit trial number + tDCS condition:exit trial number \| participant) |

MSIT = Multi-source interference task, RT = response time, NFC = Need for Cognition.
Regression equations correspond to the code deposited on OSF (available upon final publication). Regressions included maximal random effects structure that yielded model convergence.

### Additional modeling details

As described in the main manuscript, we estimated a hierarchical Bayesian logistic mixed-effects model based on the MVT (Charnov, 1976) to predict participants' stay (i.e., harvest) and exit (i.e., travel) decisions, and to derive participant-specific estimates for the changes in travel cost as due to environment effort level and stimulation condition. The approach follows the analysis by (Bustamante et al., 2023). Specifically, we calculated the probability for a participant, $i$, to stay in a low or high-effort orchard on a given trial $t$ separately for both stimulation conditions as follows:

$$P(stay)_{it} = \frac{1}{1+\exp\left(\beta_i\left(R_{e_{it}}-\rho_{it}\right)\right)} \, , \quad (1)$$

where $p$(stay) denotes the probability to stay (or harvest) a tree in a low or high-effort orchard, $\beta$ denotes the inverse temperature of the softmax function, indicating choice stochasticity with higher values corresponding to more deterministic choice behavior. $R_e$ is the expected reward of the next harvest defined as the last reward multiplied by the mean depletion rate (κ). The first harvest of a patch, being forced, was excluded from analysis:

$$R_{e_{it}} = r_{t-1} * \kappa, \quad (2)$$

where $r_{t-1}$ is the average reward of the previous harvest and κ is a constant of 0.88, reflecting the mean depletion rate of rewards across harvests. $\rho$ is the average reward rate of the current block type (low or high effort) given by:

$$\rho_{it} = \frac{\sum_j^t r_j - \sum_j^t c_{it}}{\sum_j^t T_{ij}}, \quad (3)$$

where $c$ is the travel cost associated with effort defined as:

$$c_{it} = \begin{cases} c_{low_i}, & E_{it} = 0 \\ c_{low_i} + c_{high_i}, & E_{it} = 1 \end{cases} \quad (4)$$

and $T$ is a variable used to count how many previous trials were completed in this effort level (i.e., the cumulative time spent time in a block type), defined as:

$$T_{it} = \begin{cases} 1, & E_{it} = 0, \text{if } c_{it} = c_{low} \\ 1, & E_{it} = 1, \text{if } c_{it} \neq c_{low} \\ 0, & \text{otherwise} \end{cases} \quad (5)$$

In equations (4) and (5), $E$ is a dummy-coded effort level variable, which is 0 if the current trial is part of a low-effort block (i.e., congruent MSIT) and is 1 if it is part of a high-effort block (interference MSIT).

As a central part of the model, the travel cost $c$ is instantiated by two separate paramaters (see equation (3)): $c_{low}$ and $c_{high}$, where $c_{high}$ is expressed as the marginal increase in travel cost from low (represented by $c_{low}$) to high effort in the respective stimulation conditions—in other words, the effect for high effort cost represents the difference in costs between effort conditions. This parameterization allows us to capture individual biases in baseline (low-effort) exit thresholds between participants.

So far, the model specifications are identical to those described by (Bustamante et al., 2023). To additionally assess the impact of stimulation condition on foraging decisions unique to the current study, both cost parameters (i.e., $c_{low}$ and $c_{high}$) as well as the inverse temperature $\beta$ were in turn predicted as a linear combination of an intercept term and a dummy-coded stimulation condition variable, $S$ (-1 = sham, 1 = anodal), such that:

$$\begin{bmatrix} \beta_i \\ c_{low_i} \\ c_{high_i} \end{bmatrix} = \left( \begin{bmatrix} \gamma_{00} \\ \gamma_{01} \\ \gamma_{02} \end{bmatrix} + \begin{bmatrix} U_{00i} \\ U_{01i} \\ U_{02i} \end{bmatrix} \right) \cdot \begin{bmatrix} 1 \\ 1 \\ 1 \end{bmatrix} + \left( \begin{bmatrix} \gamma_{10} \\ \gamma_{11} \\ \gamma_{12} \end{bmatrix} + \begin{bmatrix} U_{10i} \\ U_{11i} \\ U_{12i} \end{bmatrix} \right) \cdot \begin{bmatrix} S_{it} \\ S_{it} \\ S_{it} \end{bmatrix}, \tag{6}$$

where $\gamma$ represents the fixed effect for each parameter and $U$ represents a participant-level deviation from $\gamma$ (i.e., a random effect). Random effects were assumed to be fully correlated, such that:

$$U \sim MVN(0, \Sigma) \tag{7}$$

and

$$\Sigma = \begin{bmatrix} \tau_{00}^2 & \omega_{01,00} & \cdots & \omega_{11,00} & \omega_{12,00} \\ \omega_{00,01} & \tau_{01}^2 & \cdots & \omega_{11,01} & \omega_{12,01} \\ \vdots & \vdots & \ddots & \vdots & \vdots \\ \omega_{00,11} & \omega_{01,11} & \cdots & \tau_{11}^2 & \omega_{12,11} \\ \omega_{00,12} & \omega_{01,12} & \cdots & \omega_{11,12} & \tau_{12}^2 \end{bmatrix}, \tag{8}$$

where $\tau$ is the random variance term, $\omega$ is the covariance between parameters' random effects, and all subscripts correspond to the effects enumerated in Equation 6.

Overall then, the model had six free fixed-effect parameters ($\gamma$s, see above), six random-effect parameters ($\tau$s), and a $6 \times 6$ covariance matrix of random effects. Priors on fixed effects were assumed to be normally distributed with a mean hyperparameter of 0 and standard deviation of 0.5, 40, 30, 0.25, 20, 15 for $\gamma$s 01 to 12 respectively. Priors on random variance were assumed to be half-normal with means 0.25, 20, 15, 0.15, 10, 5 and standard deviation of 0.25, 20, 15, 0.15, 10, 5 for $\tau$s 01 to 12 respectively. The covariance matrix Σ was defined using a Lewandowski-Kurowicka-Joe distribution with shape 1 (Lewandowski et al., 2009). Participant-level parameters and their group-level distributions were estimated using Markov Chain Monte Carlo (MCMC) sampling implemented in Stan with the CmdStanR package (4,000 samples, 2,000 warm-up samples, 4 chains; Stan, 2021).

For model-agnostic analyses, we used linear mixed-effects regression to predict log transformed expected reward.

$$R_{e_{it}} = \frac{r_{t-1} + r_{t-2} * \kappa}{2}, \qquad (8)$$

The first harvest of a patch was excluded from analysis, and on the second harvest of a patch we used the last reward multiplied by the depletion rate.

<u>Model comparison</u>

To provide additional evidence for the presence of a stimulation effect on computational parameters in the EFT model, we conducted a formal model comparison using approximated leave-one-out cross-validation information criterion (LOO-IC, Vehtari et al., 2020) for (1) a null model (the standard EFT model without stimulation or session effects, as described in Bustamante et al. 2023, Experiment 2), (2) a model that assumes only stimulation effects (but no session effects, this is the model we reported), and (3) a model that assumes both stimulation and session effects—in other words, a “forward-selection” Bayesian model comparison approach. All three models converged (R-hat diagnostic statistic < 1.010 for all parameters of the null model, R-hat < 1.005 for the stimulation effect model, R-hat < 1.004 for the stimulation and session effect model).

In doing so, we found that the stimulation effect model yielded the best fit to our data. Numerically, LOO-IC under the stimulation effect model (19166.5) was superior to the LOO-IC under the null model (21486.7), indicating that the improvements in log-likelihood from including the session effects were justified by the added complexity of stimulation effects. The (absolute) difference in elpd (expected pointwise log posterior density) between the null and stimulation models was 1,160.1, with a standard error of 39.1. Using the standard 1.96 cutoff rule (1.96 * 39.1), the difference between elpd would have to be at least 76.64 to be considered "significant". As 1,160.1 > 76.64, we conclude that the stimulation model is significantly better fit than the null model (Vehtari et al., 2020; Devine et al., 2023). The LOO-IC under the stimulation and session model was the highest (39648.0), so the addition of session effects was not justified.

### Supplementary results

#### OSPAN effects

**Table S3.** Results from a mixed-effects regression model predicting exit thresholds by stimulation condition, effort level, and OSPAN performance.

| Predictor | β (SEM) | *df* | *t*-value | *p*-value | |
|---|---|---|---|---|---|
| Intercept | 6.84 (0.36) | 43.99 | 19.24 | <.001 | *** |
| Effort | -0.30 (0.06) | 41.64 | -4.76 | <.001 | *** |
| tDCS condition | -0.32 (0.54) | 43.72 | -0.60 | .553 | |
| OSPAN total | -0.03 (0.34) | 73.74 | -0.08 | .937 | |
| Effort × tDCS condition | 0.21 (0.11) | 40.19 | 1.91 | .064 | |
| Effort × OSPAN total | -0.07 (0.06) | 65.83 | -1.09 | .281 | |
| tDCS condition × OSPAN total | -0.97 (0.57) | 54.21 | -1.69 | .096 | |
| Effort × tDCS condition × OSPAN total | -0.04 (0.11) | 53.93 | -0.39 | .700 | |

*** p < .001, ** p < .01, * p < .05

#### Time-on-task effects

**Table S4.** Results from a mixed-effects regression model predicting exit thresholds by stimulation condition, effort level, and exit trial number within blocks.

| Predictor | β (SEM) | *df* | *t*-value | *p*-value | |
|---|---|---|---|---|---|
| Intercept | 6.81 (0.34) | 45.02 | 19.73 | <.001 | *** |
| Effort | -0.29 (0.06) | 42.98 | -4.49 | <.001 | *** |

| | | | | | |
|---|---|---|---|---|---|
| tDCS condition | -0.33 (0.56) | 44.98 | -0.59 | .560 | |
| Block exit trial number | -0.03 (0.03) | 39.69 | -1.01 | .318 | |
| Effort × tDCS condition | 0.19 (0.08) | 3486.40 | 2.29 | .022 | * |
| Effort × Block exit trial number | 0.01 (0.06) | 42.42 | 0.05 | .964 | |
| tDCS condition × Block exit trial number | -0.04 (0.06) | 36.98 | -0.59 | .559 | |
| Effort × tDCS condition × Block exit trial number | 0.07 (0.09) | 3342.12 | 0.77 | .443 | |

*** $p < .001$, ** $p < .01$, * $p < .05$
